# Haplotype of the astrocytic water channel AQP4 modulates slow wave energy in human NREM sleep

**DOI:** 10.1101/2019.12.23.886952

**Authors:** Sara Marie Ulv Larsen, Hans-Peter Landolt, Wolfgang Berger, Maiken Nedergaard, Gitte Moos Knudsen, Sebastian Camillo Holst

## Abstract

Cerebrospinal fluid (CSF) flow through the brain parenchyma is facilitated by the astrocytic water channel aquaporin 4 (AQP4). Homeostatically regulated electroencephalographic (EEG) slow waves are a hallmark of deep non-rapid-eye-movement (NREM) sleep and have been implicated in the regulation of parenchymal CSF flow and brain clearance. The human *AQP4* gene harbors several single nucleotide polymorphisms (SNPs) associated with AQP4 expression, brain-water homeostasis and neurodegenerative diseases. To date, their role in sleep-wake regulation is unknown. To investigate whether functional variants in *AQP4* modulate human sleep, nocturnal EEG-recordings and cognitive performance were investigated in 123 healthy participants genotyped for a common eight-SNP *AQP4*-haplotype. We show that this *AQP4*-haplotype is associated with distinct modulations of NREM slow wave energy, strongest in early sleep and mirrored by changes in sleepiness and reaction times during extended wakefulness. The study provides the first human evidence for a link between AQP4, deep NREM sleep and cognitive consequences of prolonged wakefulness.

## Introduction

Glial-dependent cerebrospinal fluid (CSF) flow through the brain parenchyma, by some termed the glymphatic system, facilitates a circulation of nutrients and removal of waste by generating a convective flow of CSF and interstitial fluid (1,2). Evidence suggests that the pathway relies on three main processes: Firstly, bulk flow of CSF through perivascular spaces is generated by arterial pulsations from the heartbeat and respiration (3,4). Secondly, movements of CSF from the perivascular space into the brain parenchyma relies on the water channel aquaporin 4 (AQP4), which is highly expressed on astrocytic vascular endfeet (1,5). Mice lacking AQP4 show a strong reduction in parenchymal CSF influx (5,6) and increased interstitial beta-amyloid depositions (7), which is ameliorated by sleep deprivation (8). Thirdly, the inward flow of CSF through AQP4 channels mainly occurs during non-rapid eye-movement (NREM) sleep (9) and in preclinical studies, glymphatic flow is known to be positively correlated with slow wave production (10).

Emerging data support the existence of sleep driven CSF movements and clearance in the human brain. Increased levels of intracerebral tau and ß-amyloid have been observed in healthy adults after sleep loss (11,12), and recently sleep driven parenchymal CSF pulsations in the fourth ventricle were demonstrated in the human brain (13), providing the first evidence for human glymphatic mechanisms. To date, however, no studies have described the link between *AQP4* and human sleep-wake regulation or investigated whether genetic modulation of AQP4 may have restorative effects on cognitive functions after sleep loss.

The gene encoding AQP4 is located on chromosome 18 (18q11.2-q12.1) (14) (Figure 1). Several thousand single nucleotide polymorphisms (SNPs) in the non-coding regions of *AQP4* have so far been identified and their function(s) in the normal and diseased human brain is an active area of research. Human *AQP4* SNPs have been shown to impair cellular water permeability and water homeostasis in vitro (15). Moreover, an array of studies have associated human *AQP4* SNPs with neurological disorders including Alzheimer’s disease progression (16), vascular depression phenotype (17), leukariosis (18), outcome after traumatic head injury (19), edema formation (20,21) and the risk of stroke (22). These findings suggest a link between AQP4 and the development of brain diseases associated with waste deposition and fluid movements. Recently, a single variant within *AQP4* was associated with a 15-20% change in AQP4 expression (23).

**Figure 1.**
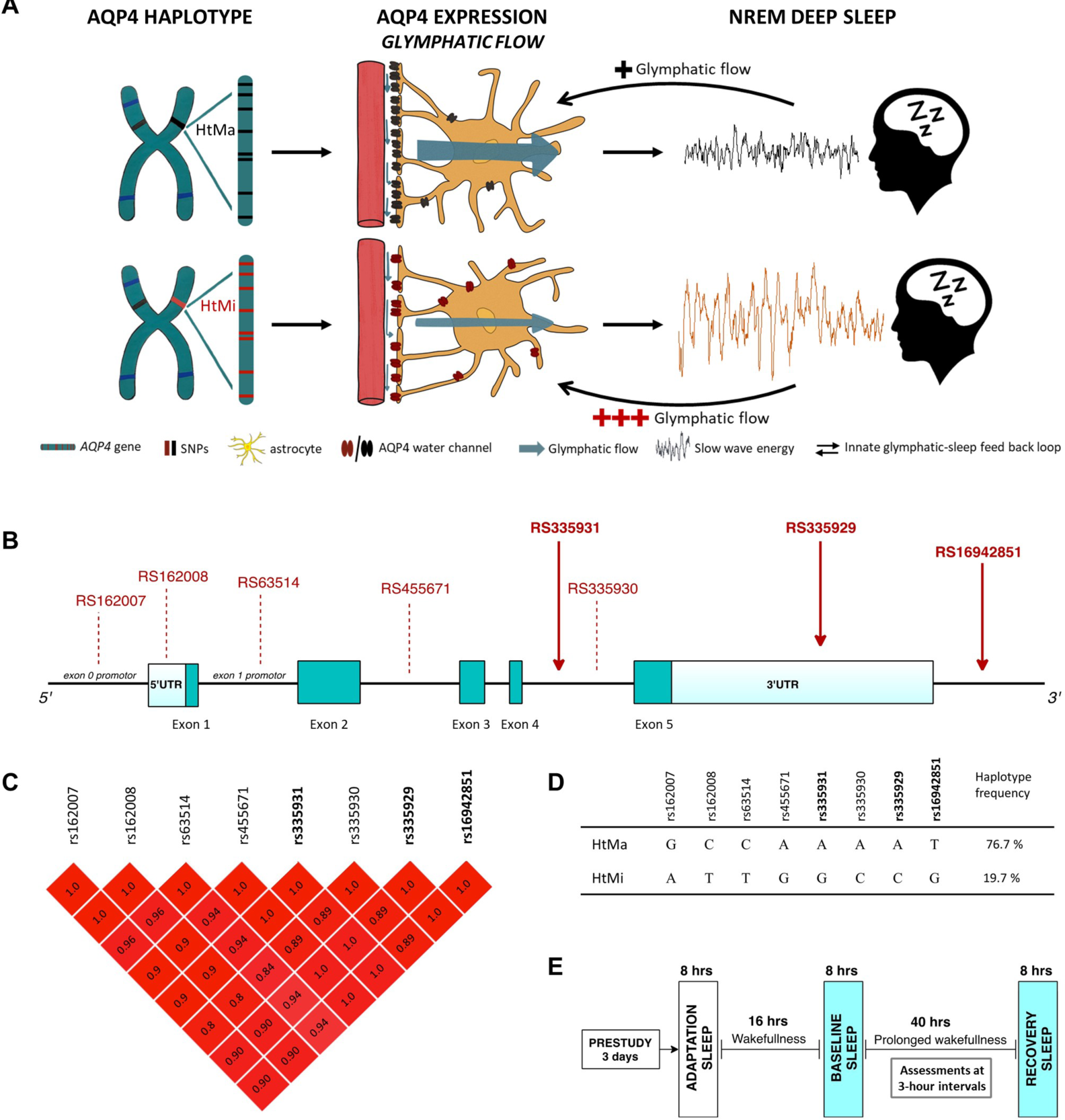
Hypothesized role of the human *AQP4* haplotype investigated in a controlled sleep deprivation study. (**A**) Based on the genetic modulation of AQP4 protein expression (23), we hypothesized that the high AQP4 expressing variant (HtMa; black) presents with improved glymphatic flow compared to the low AQP4 expressing HtMi variant (red). Assuming that NREM slow waves are the endogenous regulator of glymphatic flow, the HtMa variant require less slow wave energy (SWE) to initiate glymphatic flow than the HtMi variant, establishing an innate glymphatic-sleep feedback loop. **(B)** Physical map of the AQP4 gene and the location of the 8 haplotype SNPs. The three SNPs genotyped in this sample are marked with full red arrows. Dark green blocks: coding exons, Light blue blocks: 5′- and 3′-untranslated regions. **(C)** Linkage disequilibrium (LD) block among the *AQP4* SNPs in the investigated haplotype. The pairwise LD coefficients (r^2^) of SNPs in the LD block are color-scaled in red tones with dark red indicating perfect LD (r^2^ = 1). **(D)** Table shows bases at the 8 different SNP locations in the AQP4-gene for the two haplotypes HtMa (76.7%) and HtMi (19.7%), and their respective frequencies in the CEU and TSI populations, representative of the investigated Swiss cohort. 3.7% of CEU and TSI are predicted to be carriers of rare haplotype variants (42). Frequencies in investigated study population were close to the prediction (HtMa: 75.7%; HtMi: 23.5; others: 0.7%). **(E)** Visualization of study design common for all subjects included from six separate studies. After a minimum three-day inclusion period with monitored bedtimes and no caffeine intake, all study participants underwent an adaptation night in the laboratory before baseline sleep, 40 hours prolonged wakefulness and a recovery night, adding up to more than 1950 hours (123 × 2 × 8h) of included sleep EEG recordings. Subjective sleepiness ratings and the ~10 min psychomotor vigilance test (PVT) were performed at three-hour intervals.

Here, we aimed to investigate the role of AQP4 in human sleep-wake regulation. We hypothesized that if NREM slow waves are the endogenous regulator of CSF brain pulsations then a reduced expression of AQP4 should be associated with a compensatory increase in deep NREM sleep (Figure 1A). To investigate this association, a haplotype spanning *AQP4* was genotyped in a sample of 123 healthy participants from controlled sleep deprivation studies. Data from all-night electroencephalographic (EEG) recordings in baseline- and recovery-sleep, as well as measurements of subjective sleepiness and global alertness throughout 40 hours of prolonged wakefulness were analyzed (Figure 1E).

## Results and discussion

### The AQP4-gene harbors an eight-SNP haplotype associated with AQP4 expression

Initial examination of SNPs in the *AQP4*-gene revealed a conserved haplotype spanning the entire gene with two common variants (Figure 1B). This haplotype consists of eight SNPs, including rs335929 implicated in cognitive decline in Alzheimer’s patients (16) and rs162008 demonstrated to reduce AQP4 expression (23). Based on overall variance in EEG slow wave energy (SWE; 0.75 – 4.5 Hz) and the a priori aim to detect an effect size of at least 5%, power analysis suggests a required sample size of 78 (39 per group) as sufficient (see methods). The *AQP4*-haplotypes were compared by means of dominant analysis. Fifty-two subjects were carriers of the minor allele (HtMi) and 71 individuals were homozygous for the major allele (HtMa). The two groups did not differ in demographic characteristics (Table S1 & Table S2), presented with similar sleep architecture and had a normal response to sleep deprivation (Table S3).

### AQP4-haplotype modulates slow waves in NREM sleep

The sleep EEG is genetically determined, with NREM sleep exhibiting up to 90% heritability (24), making it one of the most hereditary human traits described. To investigate whether the *AQP4*-haplotype modulates homeostatic sleep-wake regulation, EEG energy in predefined frequency bands in NREM sleep in baseline- and recovery nights and the evolution of subjective sleepiness as well as cognitive performance measures were compared between the *AQP4*-haplotypes by a fixed sequence procedure (25) (see Methods). EEG SWE, which is a combined measure of sleep intensity and duration, and one of the best validated markers of sleep propensity in humans (26), was defined as the primary outcome variable. EEG quantification revealed that the HtMi-carriers produced more SWE than the HtMa homozygotes (‘genotype’: P < 0.03; Figure 2A). This effect was not observed in the spindle range or any other frequency band (data not shown), nor in REM sleep (Figure S1). The demonstrated *AQP4*-haplotype modulation of SWE documents an association between the intensity of deep NREM sleep and the expression of the AQP4-water channel, a relationship that may be central for CSF driven brain pulsations. Given how recent preclinical evidence show that the intensity of slow waves is directly linked to glymphatic influx (10), and that the complete removal of AQP4 in mice results in brain impairments after sleep deprivation (8), this suggest an important role of AQP4-mediated clearance during sleep. The results match our initial hypothesis suggesting that in an attempt to compensate for a reduced AQP4 expression (23), *AQP4* HtMi-carriers have a stronger parenchymal CSF flow and increased SWE in NREM sleep (Figure 1A). Interestingly, the *AQP4*-haplotype modulation was similar in baseline- and recovery nights (‘genotype x night’: F_1,120_ = 0.37; P > 0.55; η_p_^2^ = 0.31%), proposing that the *AQP4* modulation is present both under normal sleep conditions and following the sleep homeostatic challenge.

**Figure 2.**
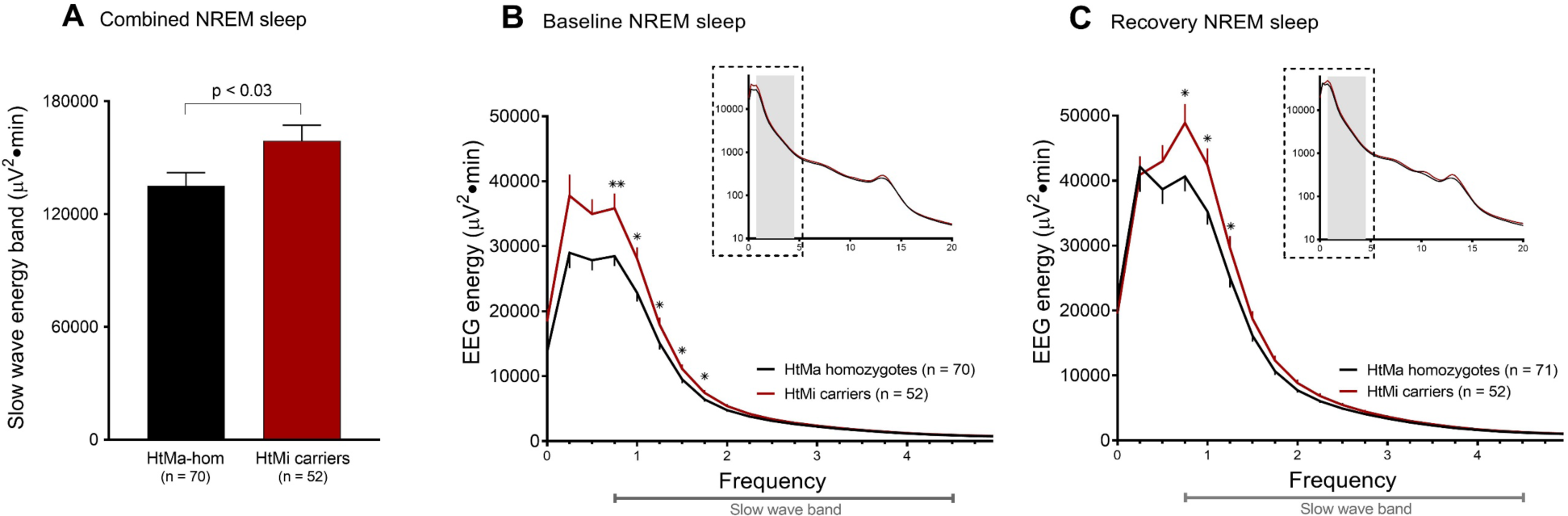
*AQP4* haplotype modulates EEG energy in the slow wave range. Comparison of EEG-energy (EEG-power x time) across baseline and recovery nights in the slow wave range (0.75-4.5 Hz) within the AQP4 haplotype variants HtMa homozygotes (black) and HtMi-carriers (red). To minimize false positive results, EEG data was analyzed by a hypothesis driven fixed sequence procedure that only revealed significant effects of AQP4 in the whole night slow wave band, which was significantly increased in the *AQP4* HtMi carrier group when compared to HtMa homozygotes (A; ‘genotype’: F_1,121_ = 5.0; P < 0.03; η_p_^2^ = 3.95%). The effect that was similar in baseline (B) and recovery (C) conditions and confined to the 0.75 – 2 Hz band. Inserts (B & C) represent full NREM sleep spectra for 0 - 20 Hz on log10 scale with grey shading indicating the slow wave band. Data represents means ± SEM. Asterix: by-bin unpaired two-tailed t-tests; * = P<0.05, ** = P<0.01.

To localize the AQP4-dependent effect in the slow wave range, bin-wise frequency analysis was performed and revealed significantly higher energy in the 0.75 – 2 Hz range for the HtMi carriers than the HtMa homozygotes (Figure 2 B-C).

### EEG markers of sleep homeostasis in NREM sleep are modulated by the AQP4-haplotype

To investigate whether the *AQP4-*haplotype modulates the well-known homeostatic decline of slow waves across sleep, EEG SWE was quantified across the first four NREM sleep episodes in baseline- and recovery nights. Consistent with the all-night sleep EEG analysis, a main effect of the *AQP4*-haplotype was observed (‘haplotype’: P < 0.03; Figure 3A - B), confirming the overall increased SWE levels in the HtMi-carrier group. Importantly however, the haplotype effect was not constant across the night. Rather, *AQP4* HtMi-carriers showed increased EEG energy mainly in the early part of the night (‘haplotype x NREM episode’: P < 0.05; Figure 3A - B) where sleep pressure is highest. This finding may suggest an *AQP4*-dependent homeostatic modulation of NREM sleep. The effect in the first NREM episode was masked by a small delay in REM onset (8.9 ± 4.3 min) in the HtMa-homozygotes compared to the HtMi-carrier group (‘haplotype’: F_1,121_ = 4.3; P < 0.05; η_p_^2^ = 3.41%; Table S3), making the AQP4-haplotype modulation clearer when the first two NREM episodes were combined (Figure 3 - inserts). Because the data show increased SWE in the low AQP4-expressing HtMi-carriers, our findings support the presence of an innate glymphatic-sleep feedback loop (Figure 1A), and suggests deep NREM sleep, not AQP4, as the main regulator of glymphatic flow.

**Figure 3.**
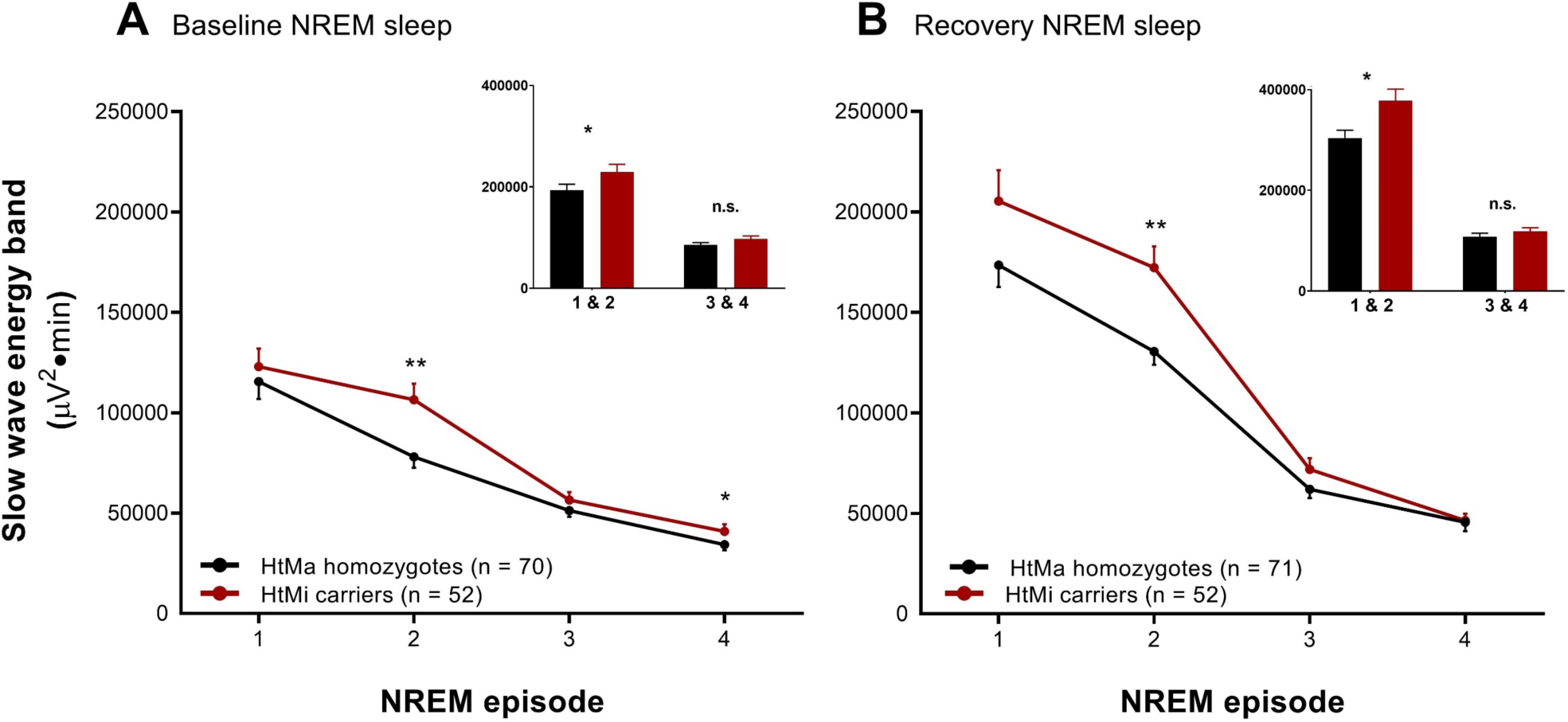
Time course of EEG slow wave production is modulated by the *AQP4* haplotype. To probe the role of AQP4 on sleep-wake regulation, slow wave energy (SWE) across the first four NREM episodes (A & B) was investigated. The data confirmed the previously detected overall increase in SWE in the *AQP4* HtMi carriers (red) compared to the HtMa homozygote (black) group (‘haplotype’: F_1,121_ = 5.39; P < 0.03, η_p_^2^ = 4.26%). Moreover, a significant AQP4-haplotype modulation across the first four NREM episodes was observed (‘haplotype x NREM episode’: F_3,843_ = 2.65; P < 0.05; η_p_^2^ = 0.93%), an effect that was strongest in the second NREM episode. AQP4 HtMi carriers were found to have increased SWE mainly in the early part of the night (figure inserts),). Spectral energy values of the slow wave band (0.75 - 4.5 Hz) in NREM sleep episodes 1-4 and the two part of the night (early:1&2, late:3&4) are plotted for the two haplotype groups for both baseline- and recovery sleep. Data represents means ± SEM. Asterix: Unpaired two-tailed t-tests; * = P < 0.05, ** = P < 0.01.

### The AQP4-haplotype modulates subjective and objective responses to prolonged wakefulness

To probe whether the *AQP4*-haplotype modulation of SWE has cognitive consequences, we investigated psychomotor vigilance performance (PVT) and subjective sleepiness (Stanford sleepiness scale, SSS) across prolonged wakefulness in the two genetic groups (see Figure 1E). Subjective sleepiness ratings revealed that HtMi-carriers coped slightly better with sleep deprivation than HtMa homozygotes and showed a smaller increase in sleepiness ratings from day 1 to day 2 (‘haplotype x day’: P < 0.04; Figure 4A). Importantly, median response speed on the PVT mirrored the effects on subjective sleepiness with the HtMi carriers reducing their speed slightly less than HtMa homozygotes (‘haplotype x day’: P < 0.04; Figure 4B), despite comparable speeds on day 1. Effects for lapses of attention were visually similar yet did not reach significance (Figure 4C). These data suggest that alterations in AQP4-dependent parenchymal CSF flow also have cognitive consequences, unveiled during sleep deprivation. Our data extends the recently described connection between slow waves, neuronal activation and CSF flow in the 4th ventricle (13), by showing that AQP4 haplotype in turn modulate NREM slow wave energy and the restorative effect of sleep on cognitive functions. These observations may indirectly suggest that AQP4 dependent fluid flow within the neuropil is regulated by EEG slow waves during NREM sleep.

**Figure 4.**
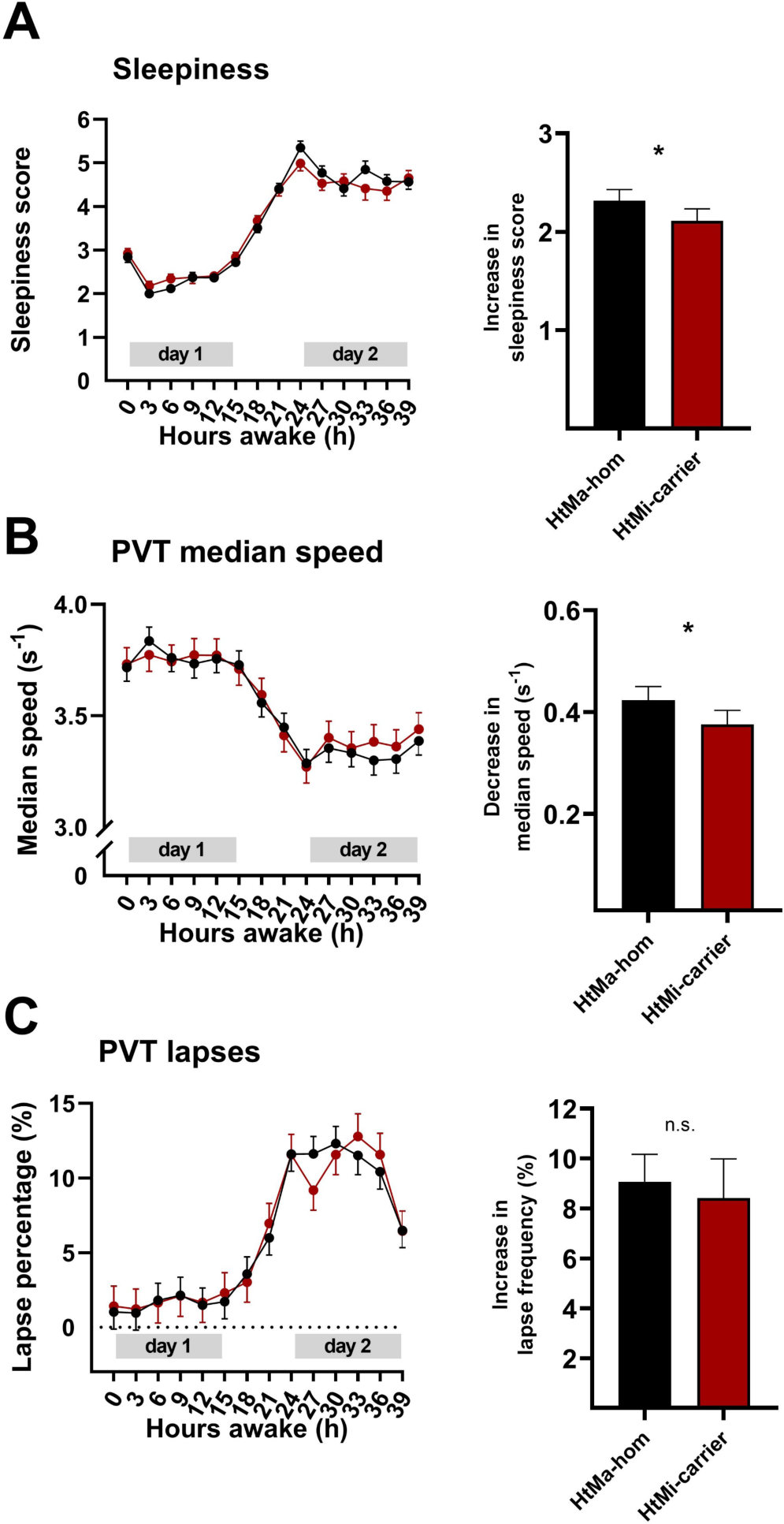
Objective and subjective measures of sleep deprivation is affected by the *AQP4* haplotype. Subjective sleepiness ratings on the Stanford sleepiness scale (SSS) and objective alertness measures by the psychomotor vigilance test (PVT). SSS scores **(A)**, median speed **(B)** and attention lapses **(C)** on the PVT were quantified at three-hour intervals across the 40 hours of prolonged wakefulness. Three-way linear mixed model analysis revealed strong differences between day 1 and day 2 (‘day’: F_all_ > 442, p < 0.0001) and moderate modulations of clock time (F_all_ >4.6, p < 0.0005) and by the ‘day x clock time’ interaction (F_all_ > 3.5, p < 0.004) in all three measures. Comparison of subjective sleepiness between the 71 HtMa homozygotes (black) and the 52 HtMi carriers (red) of the AQP4 haplotype revealed that the HtMi group coped slightly better with prolonged wakefulness than the HtMa homozygotes (‘haplotype x day’: F_1,364_ = 4.5; P < 0.04; η_p_^2^ = 1.22%; Panel A right). Intriguingly, the AQP4 dependent modulation of sleepiness was mirrored by a similar effect on PVT median speed performance among the 60 HtMa homozygotes and 44 HtMi carriers that were tested (‘haplotype x day’: F_1,1041_ = 4.7; P < 0.04; η_p_^2^ = 0.45%; Panel B right), an effect similar yet not significant for lapses of attention (‘haplotype x day’: F_1,1041_ = 0.1, p > 0.78, Panel C right). Data represents estimated means ± SEM. Asterix represent significant p-values (< 0.05) from the corresponding ‘haplotype x day’ interaction.

### Conclusion

Our data highlights that subjects carrying the low AQP4-expressing HtMi variant have enhanced SWE mainly in early NREM episodes and cope slightly better with prolonged wakefulness. Given that SNPs associated with the HtMi variant also affect the cognitive decline in Alzheimer’s patients (16), our data provides novel evidence for the existence of sleep dependent AQP4-driven CSF pulsations in the human brain consistent with proposed glymphatic mechanisms. It also supports the hypothesis that sleep slow waves are part of the regulatory machinery of parenchymal CSF flow. Further studies investigating the AQP4-haplotype and its association to sleep-associated brain functions are warranted.

## Methods

### SNPs of the AQP4 gene and haplotype analysis

To investigate genetic modulations of AQP4, common variants in the *AQP4*-gene were explored using the dbSNP database build 152 (https://www.ncbi.nlm.nih.gov/snp/). The initial search in the 1000 genome project database within dbSNP revealed 32 SNP’s with a global minor allele frequency (MAF) above 5%. Only 16 of these were common (MAF above 20%) in the European population data representative of the investigated Swiss cohort (CEU and TSI populations). Linkage disequlibrium (LD) analysis showed that eight of the 16 SNPs (rs162007, rs162008, rs63514, rs455671, rs335931, rs335930, rs335929, and rs16942851) form a distinct haplotype with SNPs in high LD (r^2^ > 0.8) (Figure 1C). Further LD analysis of the haploblock revealed that these eight SNP’s form two common variants of the haplotype: a major haplotype (HtMa) with a 76,7% incidence and a minor haplotype (HtMi) with a 19.7% incidence (for further details see Figure 1). Based on the 1000 genome database, this haplotype is widely detected across the European, American, east and south Asian populations.

### Genotyping of APQ4 SNPs

Genomic DNA extracted from 3 ml fresh EDTA-blood (wizard^R^ Genomic DNA purification Kit, Promega, Madison, WI) was used for genotyping. The rs335931, rs335929 and rs16942851 polymorphisms of *AQP4* were chosen as tag-SNP’s to represent the haplotype and were determined using Taqman® SNP genotyping Assay (Life Technologies Europe B.V.; see also Table S2). Allelic discrimination analysis was performed with SDS v2.2.2 software (applied Biosystems, Foster City, CA, USA.) All genotypes were replicated at least once for independent confirmation.

The MAFs of the genotyped variants were in accordance with MAFs predicted by the dbSNP database (Table S2). All three SNPs were in Hardy-Weinberger equilibrium. Pairwise LD coefficients (r^2^) were calculated between rs335929, rs16942851 and rs335931 confirming high linkage disequilibrium (r^2^ > 0,95). The two haplotypes and allele frequencies are shown in Figure 1.

Due to the well-established association between Alzheimer’s disease risk and Apolipoprotein E (APOE) genotype (27), we checked the distribution of APOE genotypes (rs429358 and rs7412) among the AQP4 haplotype groups using the same Taqman® SNP genotyping approach. The analysis revealed that the distribution was similar in HtMa-homozygytes and HtMi-carriers (p > 0.47; Table S1).

### Study population

To examine the impact of the genetic haplotype of *AQP4* on the sleep EEG, we investigated data from 134 healthy participants of six previously published sleep deprivation studies (28–33) All studies were conducted under strictly controlled conditions in the sleep lab of the Institute of Pharmacology and Toxicology at the University of Zürich, Switzerland using similar protocols and methodology (Figure 1E). Two carriers of rare haplotypes as well as 9 older participants with an age above 60 years were excluded from the analysis. The total sample thereby included 71 individuals homozygous for the major allele (HtMa/HtMa), 45 heterozygous (HtMa/HtMi) and 7 homozygous for the minor allele (HtMi/HtMi). Given the low number of minor allele homozygotes, a HtMi-carrier group (HtMi/HtMi and HtMa/HtMi alleles) was created (n = 52). Dominant analysis was performed with the aim of investigating the consequence of harboring the minor AQP4-haplotype. No difference in the distribution of the *AQP4*-haplotype between the six studies was observed (p > 0.21). In studies that included the administration of one or more treatments (28–30), only data from the placebo-arm were analyzed.

Study participants were right-handed healthy volunteers with a medical history free of neurological and psychiatric disorders. They were drug- and medication abstinent and reported being good sleepers with regular bedtimes and no shift-or nightwork. No participants passed through time zones or consumed excessive amounts of alcohol or caffeine in the two months prior to study-enrollment. Before inclusion, participants underwent a screening night in the sleep laboratory to check for undiagnosed sleep disorders or low sleep efficiency (<85%) (see Table S1 and S3).

The study protocols were approved by ethics committee of the Canton of Zurich for research on human subjects. Written consent was obtained from all participants before the experiments.

### Sleep study protocol

The six study protocols were very similar and were performed as follows: In the final 3 days leading up to the sleep studies, participants were required to keep a strict 8-hour/16-hour sleep schedule and to refrain from caffeine (coffee, tea, cola drinks, chocolate and energy drinks) and alcohol intake. Compliance with these requirements was verified by actigraphy from a wrist-activity monitor, sleep-wake diaries and determination of saliva caffeine as well as breath alcohol levels upon arrival in the sleep lab.

The sleep studies consisted of a block of four consecutive nights (see Figure 1): First and second nights served as adaptation and baseline-nights, respectively. The subjects were then kept awake for 40 hours (i.e. for two days, skipping one night of sleep) until bedtime on the fourth night, where they were given a 10h sleep opportunity for recovery. During the period of prolonged wakefulness, the participants were constantly supervised by members of the research team and engaged in studying, playing games, watching films, and occasionally taking a walk outside the laboratory.

### Polysomnographic recordings

Continuous all-night polysomnographic recordings were performed on all baseline- and recovery-nights. The EEG, electrooculogram (EOG), submental electromyogram, (EMG) and electrocardiogram (ECG) were recorded using the polysomnographic amplifiers PSA24 (Braintronics Inc., Almere, the Netherlands; n = 16) (28) and Artisan^®^ from Micromed (Mogliano Veneto, Italy; n = 107) (29–33). In the recordings obtained with the PSA24 recording system, the analogue EEG signals were conditioned by a high-pass 12/19/19 1:25:00 AMfilter (3 dB at 0.16 Hz) and a low-pass filter (3 dB at 102 Hz), sampled at 512 Hz, digitally low-pass filtered (3 dB at 49 Hz) and stored with a resolution of 128 Hz. In the recordings obtained with the Artisan^®^ recording system, analogue EEG data were conditioned with a high-pass filter (3 dB at 0.15 Hz), a low-pass filter (3 dB at 67.2 Hz), and sampled with a frequency 256 Hz. Sleep stages were visually scored in 20-s epochs according to standard criteria (34), and arousal- and movement-related artifacts were visually identified and removed. The data from the C3M2 derivation are reported. In both conditions, the analyses were restricted to the first 8 hours (480 min) after lights-off.

### EEG analyses

Four second EEG spectra (fast Fourier transform routine, Hanning Window, frequency resolution 0.25 Hz) were calculated with MATLAB (MathWorks Inc., Natick, MA), and EEG power spectra of 5 consecutive 4 second epochs were averaged and matched with the scored sleep stages.

The first four NREM episodes were defined according to current standards (35). The all-night power spectra represent the average of all artifact-free 20 s values in NREM sleep (stages 1-4) between 0 and 20 Hz. The energy spectra contain all values of spectral power multiplied by time (min) spent in NREM sleep per night (480 minutes) or in the respective NREM sleep episodes. The energy calculations factors in a more relevant quantitative interpretation of the spectra (35).

### Cognitive testing and sleepiness ratings

The psychomotor vigilance test (PVT) is a simple reaction time task implemented in e-Prime software (Psychology Software Tools Inc., Pittsburgh, PA), in which subjects are instructed to press a button as quickly as possible with their right index finger when they see a digital millisecond counter that starts to scroll in the center of the computer screen (36). Nineteen individuals were excluded from analysis because they performed a different, non-computerized version of the task, resulting in a sample size of 104 subjects for the PVT task analyses. Moreover, because a subset of participants underwent neuroimaging on the second day of sleep deprivation, some performance measures are missing on day 2. Subjects received oral instructions and performed a training session prior to study start. For each PVT trial, 100 stimuli were presented (random inter-stimulus intervals: 2 – 10 s). Two extensively validated PVT variables were quantified (37,38): ‘lapses of attention’ (defined as the percentage of trials with reaction times longer than 500 ms) and median response speed (based on inverse reaction times). Immediately prior to all PVT assessments, a validated German version of the Stanford Sleepiness Scale was administered (39). The sleepiness ratings of all 123 subjects were included in the analyses.

### Statistical analyses

All statistical analyses were performed with SAS 9.4 x64 software (SAS institute, Cary, North Carolina) and performed across nocturnal EEG data, subjective sleepiness and PVT performance measures. To approximate a normal distribution, EEG power was log-transformed prior to statistical tests. Two- and three-way mixed-model analysis of variance were performed with the between-subjects factor “genotype” (HtMa homozygotes vs. HtMi carriers) and the relevant within-subject factors: “condition” (baseline vs. recovery), “NREM sleep episode” (1-4), “frequency bin” (bin 1-81), clock time (8, 11, 14, 17, 20, 23 o’clock) and the duration of prolonged wakefulness (day 1 vs. day 2). For the nocturnal EEG data, two overall 4-way mixed model analysis were performed (for EEG power and EEG energy), which included all frequency bins, genotype, condition and study. Further analysis was only performed because a significant effect of genotypes was established. Moreover, to control for type I error caused by multiple comparison, the significance levels for these primary two overall mixed models were set to α < 0.025 (Bonferroni correction α = 0.05/2). Type I errors were further controlled by only considering significant effects relevant when the following two criteria were met: A. more than two bins in the by-bin (mixed model) analysis of variance were below α = 0.05 <u>and</u> B. the corresponding frequency band (slow wave/delta, theta, alpha, spindle, beta) was also significant at α level = 0.05. The final step implemented to control for type 1 errors was to investigate the sleep EEG bands using a hypothesis driven, fixed sequence procedure (25), where EEG bands were listed based on their relevance for sleep-wake regulation as follows: 1. Delta / slow wave activity (0.75 −4.5 Hz), 2. Spindles (12 - 15 Hz), 3. Theta (5 – 8 Hz), 4. Alpha (8 – 12 Hz) and 5. Beta (15 - 20 Hz), and tested in that order. Only if the previously tested band revealed a significant genotype modulation, statistical testing of the subsequent band was performed.

Based on the log transformed EEG SWE data, a priori power analysis revealed that to determine a 5% difference between haplotypes with a simple two-tailed t-test (α=0.05), a total sample size of 78 subjects (Cohen’s d: 0.833; SWE mean: 5 log[μV^2^min] ± 2.5%; standard deviation: 0.3 log[μV^2^min]) are required (G*Power 3.1.9.2; Die Heinrich-Heine-Universität Düsseldorf).

When a significant main-effect or interaction was discovered, appropriate paired or unpaired 2-tailed t-tests were used to localize differences within and between groups. If not stated otherwise, only significant effects or results are reported. Effect sizes (partial eta squared: η_p_^2^) were calculated from corresponding mixed model F-values and degrees of freedom. Effect sizes of 0.0099, 0.0588 and 0.1379 are considered small, moderate and large, respectively (40,41).

## Acknowledgments

We thank Dr. Julia Rétey, Dr. Sereina Bodenmann, Dr. Valerie Bachmann, Dr. Kathrin Stingelin and Dr. Susanne Weigend for their contributions in data collection and sleep scoring. Data acquisition was supported by grants of the Swiss National Science Foundation (grant numbers: 31-67060.01; 310000-120377; 3100A0-107874; 320030_135414; 320030_163439). We thank Dr. Marianna Di Chiara, Laura van Bommel, Silke Feil and Dr. Samuel Koller for their assistance with *AQP4* genotyping. The current study was sponsored by EU Marie Skłodowska-Curie fellowship #798131 and by a student fellowship from the Copenhagen University hospital, Rigshospitalet.

## SUPPLEMENTARY MATERIAL

**Supplementary figure 1.**
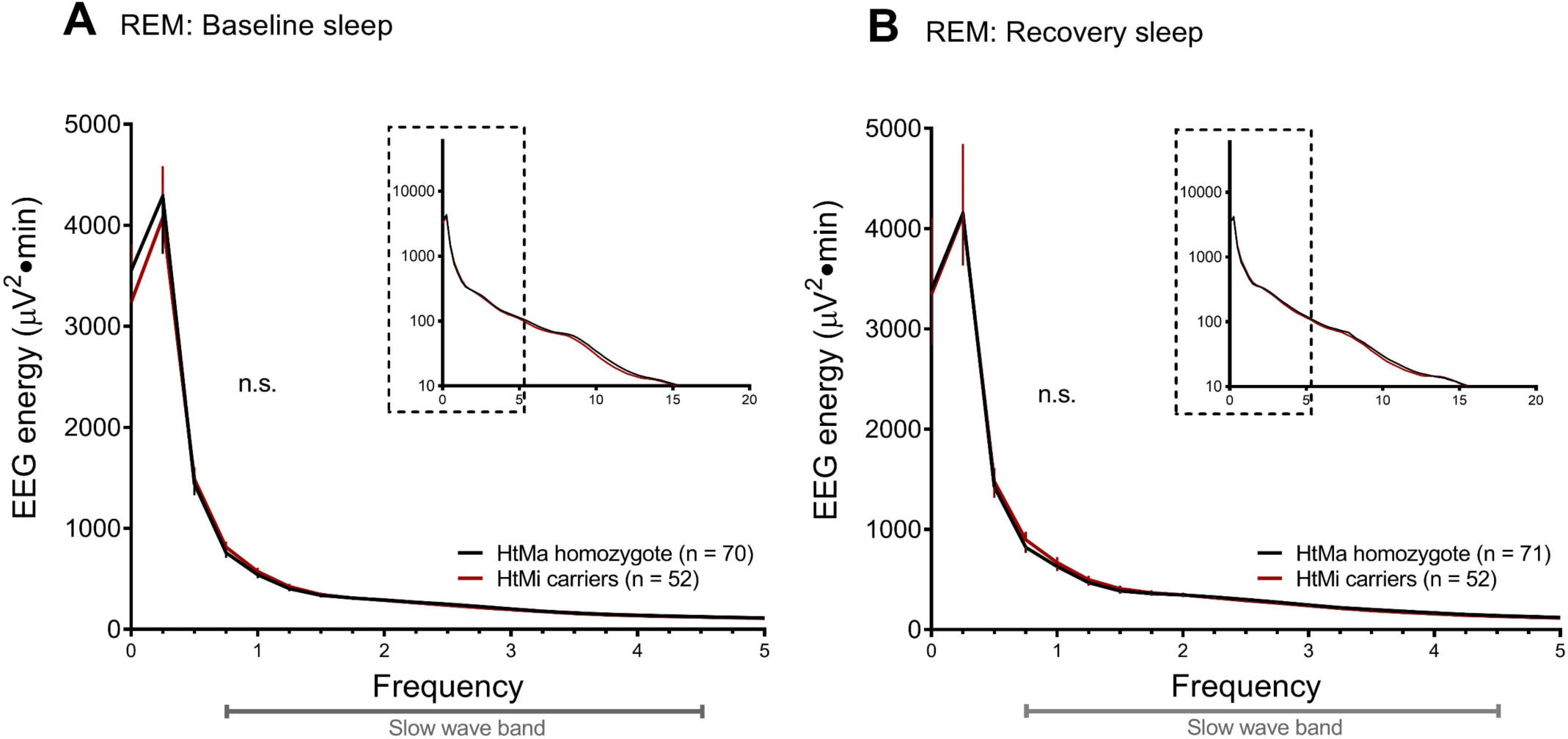
REM energy analysis for *AQP4* HtMa homozygous (black lines) and HtMi carriers (red lines) across baseline (A) nor recovery (B) nights. No significant modulation by the AQP4 haplotype was observed (‘haplotype’: F_1,21_ = 0.07; P > 0.79), confirming that the sleep EEG modulations are selective to the NREM slow wave range. Similar results were seen for power spectra analysis (data not shown). Plots represent means, error bars: SEM. Insert: Full NREM sleep spectra for 0-20 Hz on log10 scale.

**Supplementary table 1.** Demographics

|  | HtMa homozygotes | HtMi carriers | P-value |
| --- | --- | --- | --- |
| Sample size ($n_{\text{total}} = 123$) | 71 | 52 | |
| Age (years) | $24.1 \pm 2.8$ | $23.9 \pm 3.0$ | 0.69 |
| BMI ($\text{kg}/\text{cm}^2$) | $22.4 \pm 1.6$ | $22.6 \pm 2.1$ | 0.40 |
| Gender (% females) | 12,7 | 3,8 | 0.12 |
| Reported habitual sleep duration (h) | $7.3 \pm 0.7$ | $7.4 \pm 0.7$ | 0.46 |
| Trait anxiety (STAI) | $33.25 \pm 7.1$ | $35.1 \pm 8.5$ | 0.19 |
| Sleepiness (ESS) | $6.6 \pm 3.0$ | $6.9 \pm 2.9$ | 0.61 |
| Smoking (% yes) | 2,8 | 1,9 | 1 |
| Caffeine consumption (mg/day) | $108.1 \pm 100.3$ | $129.9 \pm 118.9$ | 0.27 |
| Alcohol consumption (drinks/week) | $3.2 \pm 2.7$ | $2.8 \pm 2.6$ | 0.43 |
| APOE genotype |  |  |  |
| $\epsilon 2$ carrier | 10 | 4 | |
| $\epsilon 3$ carrier | 42 | 36 | |
| $\epsilon 4$ carrier | 16 | 11 | 0.47 |

Demographic characteristics of the 123 healthy adult volunteers, who participated in one of six 40-hour sleep deprivation studies from the Zürich sleep lab where *AQP4* HtMa homozygotes and HtMi allele carriers were genotyped. No demographic differences between the two *AQP4* groups was observed. Neither was there a difference in haplotype distribution among the six studies (Fishers exact t-test, $p > 0.21$ ). German versions and validated German translations of questionnaires were used to assess lifestyle and personality traits. Questionnaires included Epworth Sleepiness Scale (ESS) (43) and State-Trait Anxiety Inventory (STAI) (44). Caffeine consumption was estimated based on average caffeine contents per serving (coffee: 100 mg, tea: 30 mg, cola drink: 40 mg [2dL], energy drink: 80 mg [2dL], chocolate: 50 mg [100 g]). The APOE genotype, known to modulate Alzheimer's disease progression, was evenly distributed among the *AQP4* haplotype groups. P-values are calculated from students two-tailed t-tests or Fishers exact test where appropriate.

**Supplementary table 2.**
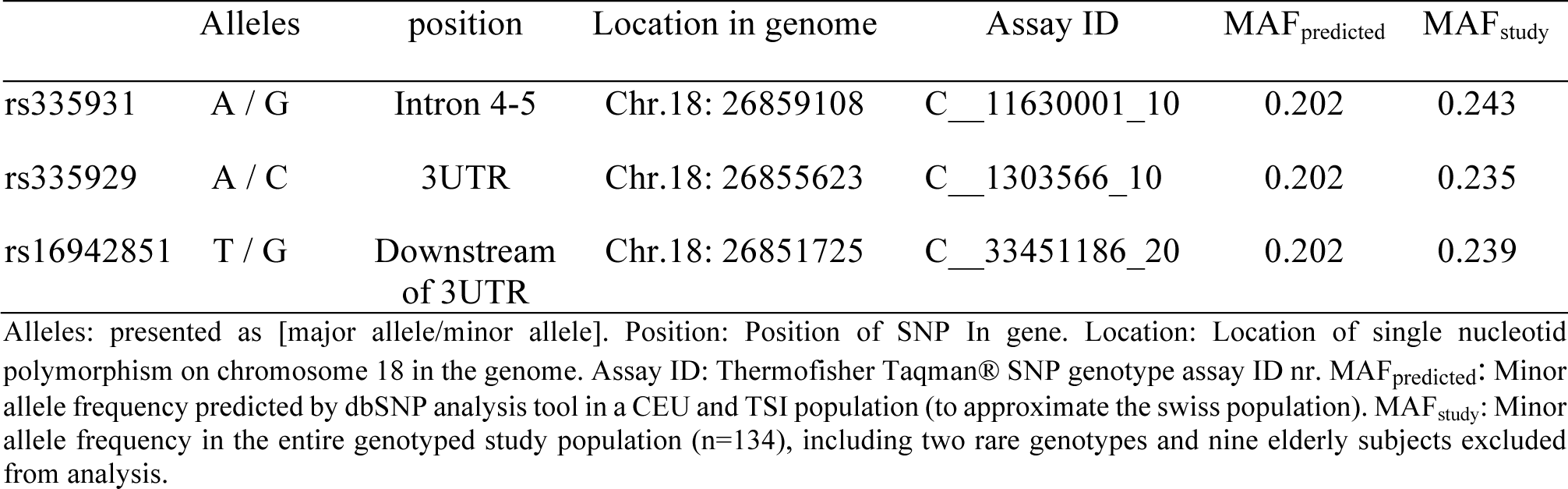
Investigated single nucleotide polymorphisms (SNPs)

**Supplementary table 3.** Visually scored sleep variables

|  | HtMa-hom (n = 71) |  | HtMi carriers (n = 52) |  | 'Genotype' |  | 'Condition' |  | 'Genotype' x 'Condition' |  |
| --- | --- | --- | --- | --- | --- | --- | --- | --- | --- | --- |
|  | Baseline | Recovery | Baseline | Recovery | F<sub>1,121</sub> | P | F<sub>1,121</sub> | P | F<sub>1,121</sub> | P |
| **Sleep efficiency (%)** | 94.0 ± 0.4 | 97.5 ± 0.2 | 94.1 ± 0.5 | 97.8 ± 0.2 | 0.3 | 0.59 | 131.0 | <0.0001 | 0.1 | 0.75 |
| **Sleep latency (min)** | 14.7 ± 1.3 | 3.5 ± 0.3 | 14.8 ± 1.6 | 3.4 ± 0.6 | 0 | 0.97 | 119.4 | <0.0001 | 0.01 | 0.92 |
| **REM sleep latency (min)** | 76.0 ± 3.5 | 81.3 ± 4.9 | 68.5 ± 2.9 | 71.0 ± 3.9 | 4.3 | 0.04 | 1.1 | 0.31 | 0.2 | 0.71 |
| **Stage N1 (min)** | 30.9 ± 2.0 | 14.8 ± 1.2 | 26.9 ± 1.9 | 12.1 ± 1.5 | 2.3 | 0.13 | 218.4 | <0.0001 | 0.4 | 0.55 |
| **Stage N2 (min)** | 215.7 ± 4.1 | 201.6 ± 4.8 | 219.9 ± 5.2 | 207.3 ± 5.6 | 0.6 | 0.45 | 25.9 | <0.0001 | 0.08 | 0.77 |
| **Stage N3 (min)** | 95.3 ± 4.1 | 149.2 ± 4.9 | 101.4 ± 5.5 | 151.2 ± 5.7 | 0.3 | 0.56 | 687.1 | <0.0001 | 1.1 | 0.29 |
| **NREM sleep (min)** | 316.4 ± 3.5 | 352.8 ± 3.1 | 323.6 ± 3.2 | 359.2 ± 3.8 | 2.5 | 0.11 | 204.5 | <0.0001 | 0.02 | 0.89 |
| **REM sleep (min)** | 109.0 ± 2.8 | 101.9 ± 3.1 | 103.0 ± 2.7 | 99.0 ± 3.6 | 1.4 | 0.25 | 6.27 | 0.014 | 0.5 | 0.49 |
| **TST (min)** | 450.9 ± 2.0 | 467.5 ± 1.0 | 451.3 ± 2.3 | 469.6 ± 0.8 | 0.5 | 0.50 | 126.5 | <0.0001 | 0.3 | 0.59 |
| **MT (min)** | 6.7 ± 0.6 | 6.7 ± 0.6 | 6.5 ± 0.6 | 5.8 ± 0.6 | 0.5 | 0.49 | 1.7 | 0.20 | 1.4 | 0.25 |
| **WASO (min)** | 7.0 ± 1.2 | 1.1 ± 0.3 | 5.9 ± 1.5 | 1.2 ± 0.4 | 0.3 | 0.61 | 35.0 | <0.0001 | 0.4 | 0.52 |
| **TIB (min)** | 479.7 ± 0 | 479.4 ± 0.6 | 479.6 ± 0.1 | 480 ± 0 | 0.5 | 0.47 | 0.01 | 0.92 | 1.0 | 0.32 |

Data on the visually scored sleep variables and their modulation by sleep deprivation and the *AQP4* haplotype. Values represent means ± SEM in baseline and recovery nights for the two haplotype groups. Analysis of the recovery nights were restricted to 480 minutes. Sleep efficiency: percentage of total sleep time per 480 min. REM sleep: rapid-eye-movement sleep. Stages N1 - N3: NREM sleep stages (N3 refers to slow wave sleep). TST: total sleep time. MT: movement time. WASO: wakefulness after sleep onset. TIB: time in bed. Sleep latency: time from lights-out to the first occurrence of N2 sleep. REM sleep latency: time from sleep onset to the first occurrence of REM sleep. F- and P-values: two-way mixed model ANOVA with factors “genotype” (HtMa-homozygotes, HtMi carriers), “condition” (baseline, recovery) and their interaction.

